# A genetically engineered porcine model of pancreatic ductal adenocarcinoma captures the biological complexity of the human disease

**DOI:** 10.64898/2026.11.16.766135

**Authors:** Tatiana Flisikowska, Laura Beltrán Sangüesa, Daniela Kalla, Christian Schneeweis, Yiran Zheng, Sandra Wiedenmann, Matthias Meier, Evelína Šťastná, Annika Eckstein, Thomas Winogrodzki, Wei Liang, Alexander Rinke Carrapeiro, Bernard Klinger, Theresa M Pauli, Kexin Wang, Katja Steiger, Christian Grätz, Michael W. Pfaffl, Rouzanna Istvanffy, Roland M Schmid, Gregers Jungersen, Ihsan Ekin Demir, Alexander Kind, Dieter Saur, Krzysztof Flisikowski, Angelika Schnieke

## Abstract

Pancreatic ductal adenocarcinoma (PDAC) is a highly aggressive cancer with limited treatment options and poor patient outcomes. The lack of clinically relevant large animal models hinders the investigation of early disease mechanisms and translational therapeutic strategies. In this study, we present a genetically engineered porcine PDAC model involving pancreas-specific activation of latent *KRAS^G12D^* alone (KC) or in combination with *TP53^R^*^167^*^H^*(KPC), developed in a midipig background. KC pigs predominantly developed progressive acinar-to-ductal metaplasia (ADM), whereas KPC pigs showed accelerated tumour progression, including PanINs, invasive carcinoma, perineural invasion, and early lymphatic dissemination. Histopathological and molecular analyses revealed pronounced intra- and inter-tumour heterogeneity, a complex stromal microenvironment, and expression of key PDAC markers. Tumours exhibited early alterations in oncogenic *KRAS* signalling, including increased dosage and differential expression of isoforms, together with dysregulation of tumour suppressor networks involving *p16* and *TP53* family members.

Single-cell transcriptomics and flow cytometric analyses demonstrated an organised and functionally active immune landscape in early lesions, characterised by infiltrating effector and regulatory T-cell populations. Porcine pancreatic cancer cell lines (PoPaCa) were established from KC and KPC tumours, recapitulating key genetic and transcriptional features of the primary lesions. Exploratory drug screening demonstrated heterogeneous responses to clinically relevant and experimental compounds. Responses to gemcitabine and paclitaxel overlapped qualitatively with those observed in human PDAC cell lines, whereas pronounced sensitivity to the GPX4 inhibitor RSL3 and partial responsiveness to trametinib identified testable therapeutic phenotypes for further validation.

Together, this genetically engineered porcine PDAC model captures early molecular, cellular, and microenvironmental features of pancreatic tumorigenesis and provides a translational platform to study disease initiation, tumour heterogeneity, immune interactions, and therapeutic response beyond the constraints of rodent models.

**Graphical abstract:** Pancreas-specific activation of mutant KRAS and TP53 in genetically engineered pigs recapitulates key features of human pancreatic ductal adenocarcinoma and provides a platform for studying tumour biology, biomarkers and therapeutic responses.

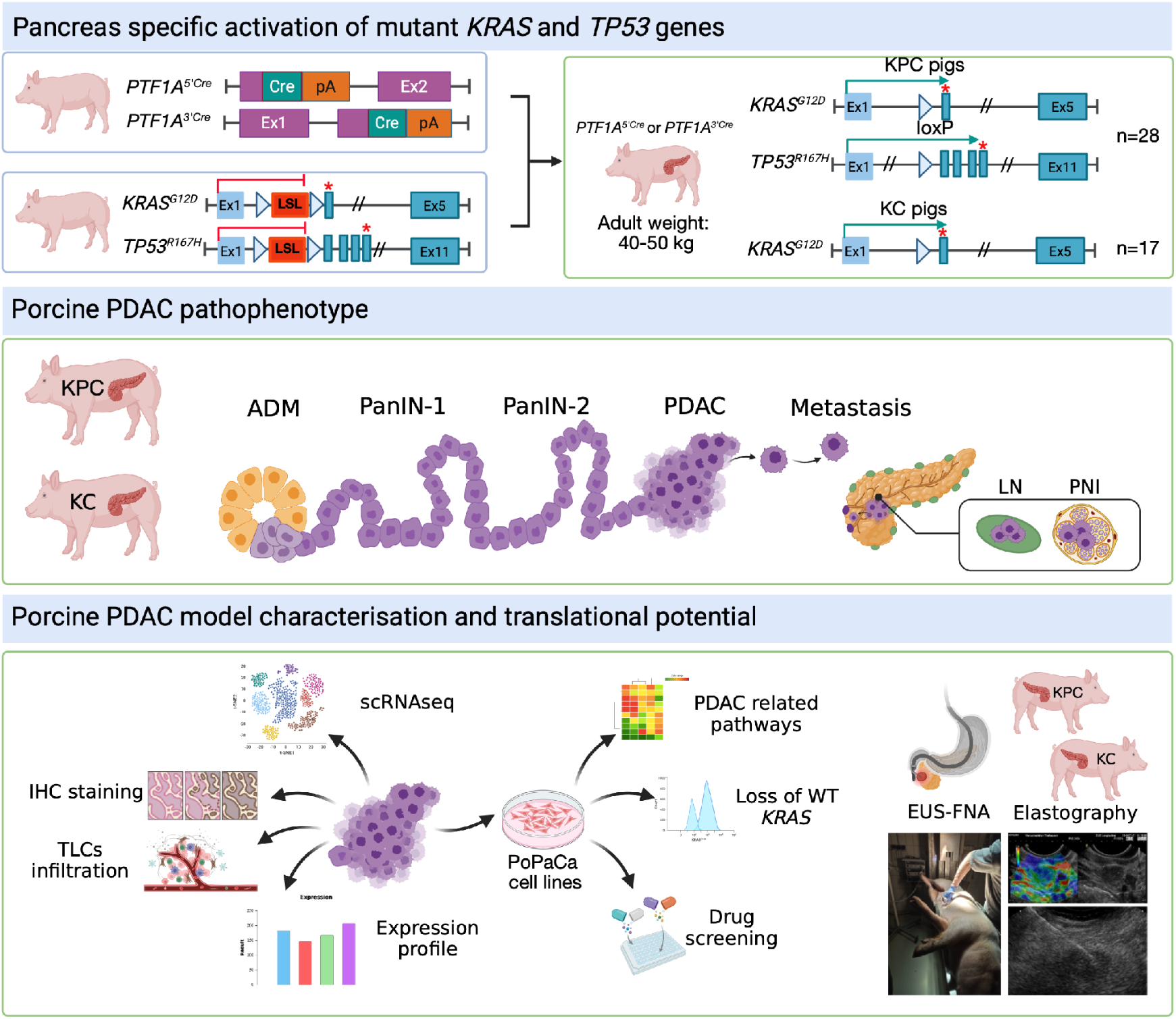

## Introduction

Pancreatic ductal adenocarcinoma (PDAC) represents a major global health issue. It exhibits one of the highest mortality rates among solid cancers, the number of patients with early-onset PDAC is increasing, and it is projected to become the second leading cause of cancer-related death in the Western world. Despite decades of research, the 5-year survival rate for PDAC has persistently remained below 10% over the past 30 years^1^ and novel systematic therapies only offered a modest life extension^2^. More recent findings have brought new hope. The development of novel KRAS inhibitors and proteolysis-targeting chimaeras (PROTACs) has shown promise in targeting mutant KRAS, a key driver of PDAC^3^. In addition, personalised RNA-based cancer vaccines are emerging as a potential therapeutic approach and have demonstrated the ability to induce robust immune responses against pancreatic tumours^4^. Despite these breakthroughs, the near-uniformly fatal disease outcome, coupled with an increasing incidence, underscores an immense medical and socioeconomic burden.

The search for better diagnosis and treatment options has motivated scientists to engineer several mouse models to mimic human PDAC. The most well-studied and commonly used are KPC mice with conditional activation of *Kras^G12D^* and *Tp53^R^*^172^*^H^*mutations in the pancreas via *Pdx1*- or *Ptf1a*-driven Cre recombinase. The KPC mice have been shown to recapitulate the initiation and development of metastatic PDAC^5^. The mouse model improved our understanding of the disease and supported essential discoveries. However, mice differ substantially in size, anatomy and physiology from humans, and longitudinal studies and complex therapeutic interventions, such as repeated tissue sampling or local ablation therapies are difficult in small animal models.

No animal model can fully replicate the complex process of cancer development in humans. Pigs, however, do have advantages and play an increasingly important role in preclinical research. They share many similarities with humans in terms of body size, anatomy, physiology and pathophysiological responses. Studies by us and others have demonstrated the suitability of pig models for both basic and translational cancer research^6–13^. Based on the clinical need, considerable efforts have been made to generate a large animal PDAC model, but so far with limited success^14–17^.

Here, we report the development and characterisation of a spontaneous porcine PDAC model that closely mimics the disease in humans and is ideally suited for translational research.

## Material and Methods

### Ethics statement

Animal experiments were approved by the Committee on Animal Health and Care of the local government body of the state of Upper Bavaria (Permission No. 55.2-2532.Vet_02-18-33) and performed according to the German Animal Welfare Act and European Union Normative for Care and Use of Experimental Animals.

### Pig housing

The different lines of midipigs (German Landrace x Minipig) were housed at the TUM Animal Research Center (ARC). Piglets were weaned at five weeks. All animals received the same diet (HEMO U 134 pellets (LikraWest)) and water *ad libitum*. Health status was continuously monitored, and experiments were terminated upon signs of deterioration of health.

### DNA extraction

DNA isolation from cells and bulk tumour samples was done with QuickExtract™ DNA Extraction Solution (Biosearch Technologies) and AllPrep DNA/RNA Mini Kit (Qiagen), respectively.

### RNA isolation

RNA was isolated from tumour bulk samples and cancer cell cultures using a Monarch Total RNA Miniprep Kit (NEB) according to the manufacturer’s protocols.

### Genotyping PCR

For pig genotyping, DNA from ear punch biopsies was analysed by PCR for *KRAS*, *TP53*, *PTF1A5’^Cre^* and *PTF1A3^’Cre^*as previously described^18^.

### RT-PCR

RT-PCR was applied to determine the expression of mutant and wild-type *KRAS* alleles in tumour samples. 200 ng of RNA was used for the cDNA synthesis using LunaScript RT Master Mix Kit (NEB) according to the manufacturer’s protocol. Diluted cDNA (1:3) was used for RT-PCR (Q5 High-Fidelity DNA polymerase, BioLabs) with *KRAS* primers in exons 2 and 3. The resulting products were visualised after 2% agarose gel electrophoresis.

### Genes expression quantification

Real-time quantitative PCR (qPCR) was performed using the Qpcrbio SyGreen Mix Lo-ROX (Biosystems) in a QuantStudio 5 Real-Time-PCR-Cycler (Thermofisher Scientific) with default thermal cycling parameters. Reactions were performed in 10 µL volume. Samples were assayed in triplicate, and the relative expression of the bulk tumour samples was normalised to *EIF2B1* expression. Expression differences were calculated using the ΔΔCt method^19^. One-way ANOVA was performed, followed by Tukey’s test for statistical evaluation using GraphPad Prism 8. The list of primers used for qPCR is shown in Supplementary Table S1.

### Recombination PCR

Excision of the loxP-STOP-loxP (LSL) cassette, inserted into intron 1 of both *KRAS* and *TP53*, was confirmed using PCR across the cassette (see Table S1 for primer sequences). PCR product was amplified by using Q5 High-Fidelity DNA polymerase (BioLabs) and the following cycling conditions: 30 s, 98 °C, then 35 cycles of 10 s, 98°C, 20 s, 60°C, 3 min, 72°C followed by 5 min at 72°C. PCR analysis of DNA extracted from healthy or tumour tissue resulted in three characteristic amplification products for either the wild type, the recombined or unrecombined allele for *KRAS* (167 bp, 203 bp and 1878 bp) or *TP53* (198 bp, 238 bp and 1520 bp).

### Determination of *KRAS* allele frequencies

Pyrosequencing was used to measure the proportion of mutant (G) and wild-type (A) *KRAS* alleles. Pyrosequencing *KRAS* allelic quantification assay was designed using PyroMark Assay Design 2.0 software (Qiagen). 500 ng genomic DNA was bisulphite-converted with the EZ DNA Methylation-Direct kit (Zymo Research, Irvine, USA) according to the manufacturer’s instructions. PCR samples were amplified using a PyroMark PCR kit (Qiagen). PCR products were sequenced using PyroMark Q48 Advanced CpG reagents on a PyroMark Q48 Autoprep instrument (Qiagen) and analysed using PyroMark Q48 software (Qiagen) to determine the relative proportions of mutant and wild-type alleles.

### Isolation of extracellular vesicles and cell-free RNA

Extracellular vesicles (EVs) were concentrated from the plasma samples by precipitation with 8% v/v 6 kDa polyethylene glycol (PEG-6000). After incubation on ice for 2 h, the precipitated EVs were pelleted by centrifugation at 1,500 g for 30 min and resuspended in 240 µL PBS. After addition of 700 µL Qiazol lysis reagent (Qiagen 79306) to the EV preparation, total cell-free RNA (cfRNA) was isolated using the miRNeasy Micro Kit (Qiagen 217084) according to the manufacturer’s instructions. Quantity and quality of the isolated cfRNA was assessed using an RNA 6000 Pico Kit on a Bioanalyzer 2100 automated electrophoresis system (Agilent Technologies 5067-1513) according to the manufacturer’s instructions.

### qPCR analysis of EV-associated KRAS and TP53 transcripts

RNA handling and all qPCR experiments were performed according to the most recent MIQE guidelines^20^. The LunaScript RT SuperMix Kit (New England Biolabs E3010) was used for reverse transcription of total cfRNA. qPCR was performed on a MIC qPCR cycler (Bio Molecular Systems) using the Luna Universal qPCR Master Mix (New England Biolabs M3003), with 40 cycles and 0.3 °C/s increments for the melting curve. No-reverse-transcriptase (NRT) controls were included in the qPCR experiments to rule out potential gDNA contaminations in the RNA isolates. Relative gene expression was calculated from the RT-qPCR results using the ΔΔCq method^19^. The Wilcoxon signed-rank test was performed to compare log2 fold change values. The list of primers is shown in Supplementary Table S1.

### Histology and immunohistochemistry

Tumours identified during necropsy were documented and evaluated by a certified veterinary pathologist. Tumour specimens for histology were fixed in 4% paraformaldehyde (Carl Roth), embedded in paraffin (FFPE) and cut into 1 μm sections. Hematoxylin and eosin (H&E) staining was carried out according to standard protocols. For immunohistochemistry (IHC), specimens were incubated with the primary antibodies at 4°C overnight, followed by incubation with secondary antibodies at room temperature for 1 hour. The immunoreaction was detected using the DAB Substrate Kit, Peroxidase (With Nikel), and stained. A list of antibodies used is shown in Supplementary Table S2.

### Porcine pancreatic cancer cells isolation

Pancreas tumour samples were collected immediately after sacrificing a pig and immersed in G solution^21^. The samples were then minced using sterile surgical scissors and digested with collagenase 4G (Collagenase NB 4G, Nordmark Biochemicals) diluted in G solution at 37°C for 30-40 minutes. The digested tissue was centrifuged at 300 x g for 5 minutes, with the deceleration speed set to a minimum. The resulting pellet was resuspended in Trypsin-EDTA and incubated for 5 minutes at room temperature. After trypsin inactivation, the mixture was filtered using a 100 µm filter and centrifuged as above. The pelleted cells were cultured in a medium consisting of Advanced DMEM/F12 (Gibco), Nu-Serum IV (Corning), bovine pituitary extract (25 µg/mL) (Sigma), Insulin-Transferrin-Selenium (0.5%) (Gibco), EGF (50 µg/mL), cholera toxin (1 mg/mL), 3,3,5 Triiodo-L-thyronine (50 µM), Dexamethasone (10 mM), Nicotinamide (1M). Established porcine pancreatic cancer cell lines (PoPaCa) were cultivated as previously described^22^

### Masson’s trichrome

To detect collagen fibres (stroma content), 1 µm FFPE tumour sections were stained using Masson’s trichrome stain kit (Polyscience).

### Flow Cytometry analysis

Lymphocytes were isolated from blood, lymph nodes (LNs) adjacent to the pancreas, and pancreatic tumours. Blood and tissue samples from a wild-type (WT) pig from the same litter were included as a control. Peripheral blood mononuclear cells (PBMCs) were purified from the whole blood using SepMate™-50 tubes (StemCell Technologies) following the manufacturer’s instructions. Lymphocytes from LNs were obtained by cutting the tissue into small pieces with sterile scissors, mashing and filtering through a 70 μm cell strainer. The tissue was washed twice in PBS + 2% FCS and filtered again through a 70 μm cell strainer to ensure a single cell suspension. For isolation of tumour infiltrating lymphocytes (TILs), a Tumor Dissociation Kit (Miltenyi Biotec) and gentleMACS™ Octo Dissociator were used according to the manufacturer’s instructions. (Miltenyi Biotec). After isolation, cells were kept in RPMI-1640 Media (Sigma-Aldrich) supplemented with 10% FCS, 0.1 mM non-essential amino acids, 1 mM sodium pyruvate, 2 mM glutamine, 100 U/mL penicillin and 100 μg/mL streptomycin (final concentrations) until further handling.

The fluorochrome-conjugated antibodies used for flow cytometric analysis are shown in Supplementary Table S2. To examine cell viability and exclude dead cells, the Fixable Viability Dye eFluor 506 kit (Thermo Fisher) was used. For immune cells stained intracellularly, 1 μg/mL phytohemagglutinin (PHA, Sigma) was used as a positive control for cytokine production. Cells were stimulated for 15 h and an additional 4 h in the presence of 1× Protein Transport Inhibitor Cocktail (Thermo Fisher Scientific). Samples for analysis of *ex vivo* cytokine production were not stimulated. The Fixation/Permeabilization Solution Kit (BD Biosciences) was used according to the manufacturer’s instructions for intracellular staining. For Foxp3 detection, cells were treated with FlowX™ Foxp3/Transcription Factor Fixation & Permeabilization Buffer Kit (Bio-techne) according to the manufacturer’s instructions, followed by Foxp3 staining. The lymphocyte samples stained only extracellularly were stained directly without prior stimulation.

Data were acquired on a BD LSR Fortessa flow cytometer (BD Bioscience) and/or Attune Nxt Flow Cytometer (Thermo Fisher) and analysed using the FlowJo Data Analysis Software version 10.8.1. Gating strategy is shown in Supplementary data.

### Isolation of immune cells from KPC pig tumor week 34 and sample preparation for scRNA-seq

After the termination of the KPC pig, one of the tumors was kept on ice for 3h until the start of the preparation. The tumour was washed with 50 ml of dPBS (Thermo Fisher Scientific) and then cut into smaller pieces of about 3 mm. To isolate the immune cells, five tumor pieces were put into a 50 ml Falcon and 15 ml of 1 µg/ml Collagenase IV (Serva) in dPBS. The pieces were kept at 37°C on a shaker with 120 rpm. The supernatant was removed after 30 and 60 min. After 120 min the supernatant was centrifuged at 300 x g for 5 min. The viability of 83% was checked with Trypan blue (Sigma Aldrich) and the cells were then in a concentration of 1.8 × 10^6^ cells/ml in 1 ml dPBS with 10% fetal bovine serum (Corning). The RNA library was generated with the Chromium Single Cell 3.1 kit v3.1 (10x Genomics), following the supplier’s instructions. The chromium controller was loaded with a target cell number of 10^4^ cells. The cDNA library was sequenced with an S2 flow cell from Illumina (NovaSeq 6000).

### RNAseq analysis

Raw sequencing data were demultiplexed, aligned to the *Sus scrofa* reference genome Sscrofa11.1^23^, filtered, and barcode/UMI counts generated using CellRanger (10x Genomics). Downstream preprocessing and analysis were performed in Python using Scanpy with default parameters unless stated otherwise. Cells with >10% mitochondrial transcripts, <1000 detected genes, or genes expressed in <3 cells were excluded. No doublets were detected using Scrublet. Data were normalized to 10⁴ counts per cell, log-transformed, and highly variable genes were selected for downstream analyses.

A neighbourhood graph was constructed using the first 15 principal components and 10 nearest neighbours. Clustering was performed with the Leiden algorithm (resolution 0.4), and visualization was generated using UMAP. Cell clusters were annotated based on known marker genes and the top 500 differentially expressed genes identified by t-test ranking. Non-significant genes (adjusted p > 0.05) were removed from the NK cell gene list.

Analyses were performed using Python 3.10.10 with Scanpy (1.9.3), Anndata (0.9.1), Leidenalg (0.9.1), Louvain (0.8.0), NumPy (1.24.3), Pandas (2.0.1), SciPy (1.10.1), Matplotlib (3.7.1), Seaborn (0.12.2), and XlsxWriter (3.1.2).

### Bulk RNA-seq analysis

Total RNA was isolated from 12 PoPaCa lines. RNA integrity was assessed using the RNA 6000 Nano Kit on a 2100 Bioanalyzer (Agilent Technologies); all samples met a minimum RNA Integrity Number (RIN) of 8. Libraries were prepared using the Illumina Stranded mRNA Library Prep Kit according to the manufacturer’s protocol. Paired-end sequencing (2 × 150 bp) was performed on an Illumina NovaSeq 6000 platform, yielded 11.8 – 18.8 million total reads per sample.

Raw reads were assessed for quality with FastQC (v0.12.1) and aligned to the *Sus scrofa* reference genome Sscrofa11.1^21^ using STAR v2.7.10. Genes with fewer than 100 counts across all samples or detected in fewer than two samples were excluded from downstream analysis. Count data were normalized using the Trimmed Mean of M-values (TMM) method implemented in edgeR. Differential expression analysis was performed using the exact test based on the negative binomial distribution in edgeR, comparing tumour-intrinsic properties and CAF content. Differentially expressed genes (DEGs) were defined as those with a |log₂ fold-change| > 1 and a Benjamini–Hochberg-adjusted *P* value (FDR) < 0.05. Gene Ontology (GO) enrichment analysis and Kyoto Encyclopedia of Genes and Genomes (KEGG) pathway analysis were performed using clusterProfiler v4.18.4, with enrichment significance defined as an adjusted *P* value < 0.05.

### High-throughput drug screening

The compound library consisted of 95 different drugs targeting diverse cancer-associated pathways and is listed in Supplementary Table S3. Drug screening was performed on two CAF-free KPC-derived porcine pancreatic cancer cell lines as previously described^24, 25^. Briefly, 2000 cells per well were seeded in 96-well plates and incubated at 37°C. After overnight incubation, cells were treated with a seven-point dilution series (7 concentrations, 3-fold dilutions, highest concentration 10 μM) plus DMSO as vehicle control and assayed 72 hours later. Viability was measured using the CellTiter-Glo® Luminescence Assay (Promega), with luminescence detected using a CLARIOStar plate reader (BMG Labtech). All assays were performed in technical duplicates, with DMSO vehicle-treated cells serving as controls. The R package GRmetrics^26, 27^ was used to generate dose-response curves.

For qualitative comparison with human PDAC cell-line responses, publicly available dose-response data were obtained from the DepMap portal website ( https://depmap.org/portal) using the harmonized CTD^2^ drug screen dataset from the Broad Institute CTD^2^ Center and originally released via the Cancer Therapeutics Response Portal (CTRP). ^28, 29^. Human PDAC cell-line dose-response curves were extracted for selected compounds also included in the porcine drug screen, namely gemcitabine, paclitaxel, RSL3 and trametinib. Because the human and porcine datasets were generated under different experimental conditions and concentration ranges, the comparison was restricted to visual qualitative assessment of dose-response profiles. Direct quantitative comparison of area-under-the-curve (AUC) values between porcine and human cell lines was therefore not performed.

## Results

### Efficient pancreas specific activation of mutant *KRAS* and *TP53*

To develop a large animal model for translational pancreatic cancer research, the previously generated pig lines carrying latent *KRAS^G12D^*^30^ and *TP53^R^*^167^*^H^* ^31^ were crossed with two Cre driver lines, which had the codon-improved Cre (iCre) inserted into either the 5’ (*PTF1A^5’Cre^*) or 3’UTR (*PTF1A^3’Cre^*) of the porcine *PTF1A* gene^18^. While the placement of iCre at the 3’UTR does not interfere with *PTF1A* expression, insertion at the 5’ position caused the inactivation of the mutant allele. Since *PTF1A* is a tumour suppressor gene^32^, the latter could affect tumorigenesis. Functionality and tissue specificity of both *PTF1A^Cre^*lines had been confirmed by breeding them to dual-fluorescent reporter pigs^18^.

Breeding of the *KRAS^G12D^* and *KRAS^G12D^/TP53^R^*^167^*^H^* with iCre driver lines resulted in 28 pigs with activated mutant *KRAS* and *TP53* (KPC) and 17 animals expressing mutant *KRAS* only (KC). For the KC pigs, eight carried the *PTF1A^5’Cre^* and nine the *PTF1A^3’Cre^*transgene; for KPC animals, 18 were transgenic for *PTF1A^5’Cre^*and 10 for *PTF1A^3’Cre^*. At one month of age, one animal of each genotype was euthanised, and organs were collected and assessed for Cre mediated excision of the Lox-Stop-Lox (LSL) cassette from the *KRAS^G12D^*and *TP53^R^*^167^*^H^*alleles. Both iCre-driver lines supported efficient excision of the LSL cassette in the pancreas. In line with the data obtained from the reporter pigs, recombination was also observed in the cerebellum for *PTF1A^5’Cre^*and *PTF1A^3’Cre^* but not in other organs (Figure 1A).

**Figure 1.**
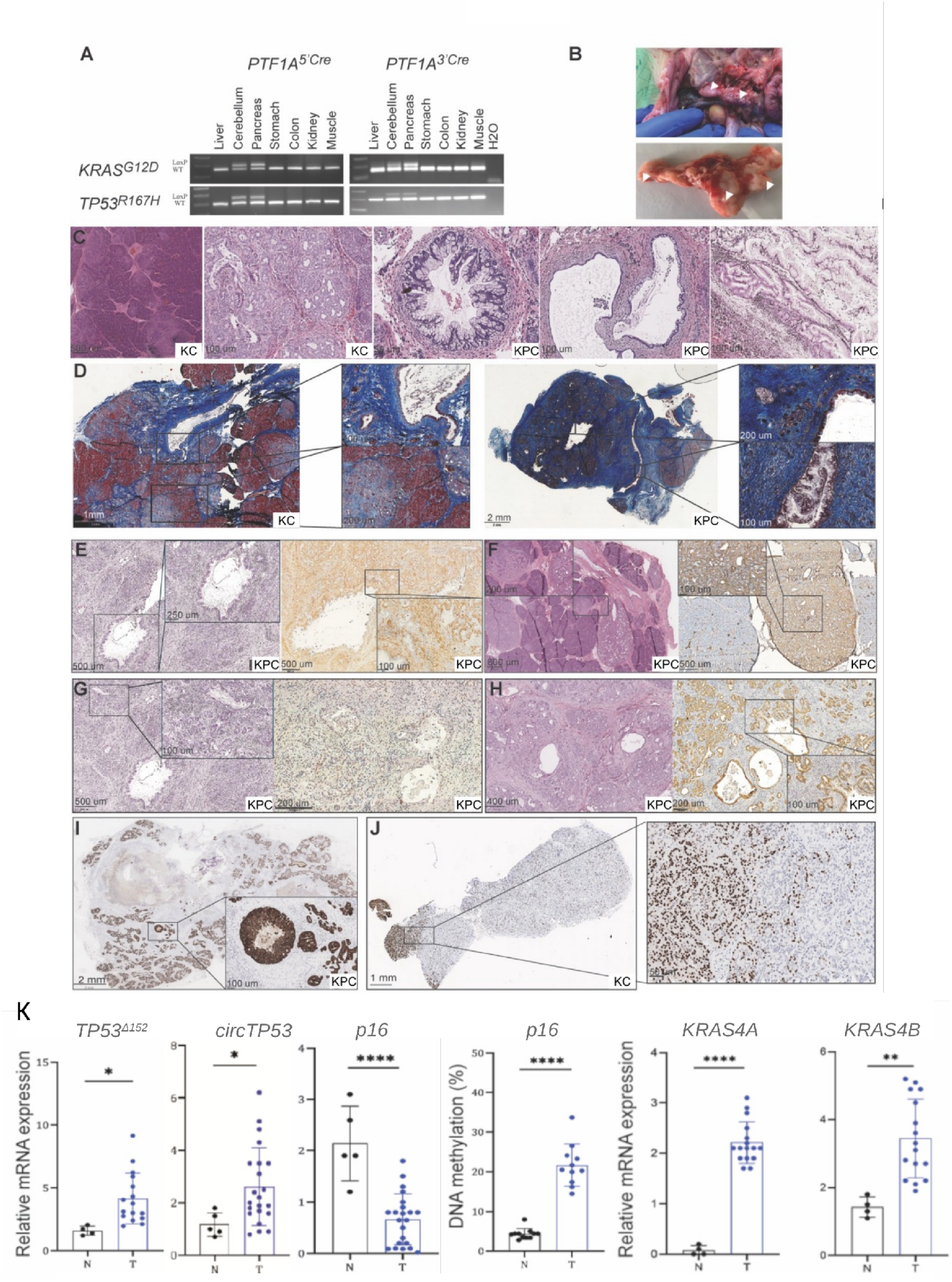
Characterisation of tumour development in the porcine PDAC model. **(A)** Gel electrophoresis image from PCR targeting Cre-mediated recombination of the *KRAS^G12D^*and *TP53^R^*^167^*^H^*alleles in different tissues from KPC pigs carrying the *PTF1A^5’Cre^*or *PTF1A^3’Cre^* driver lines. The upper band (LoxP) corresponds to Cre recombination and excision of the LSL cassette, and the lower band represents the wild-type (WT) allele. **(B)** Macroscopic appearance of pancreatic tumours in KPC pigs. **(C)** Representative H&E-stained sections showing, from left to right: focal acinar-to-ductal metaplasia (ADM), extensive ADM (KC pig), low-grade PanIN-1B (KPC pig), ducts with inflammatory infiltrates and periductal fibrosis (KPC pig), and low-grade intraductal papillary mucinous neoplasm (IPMN) (KPC pig). **(D)** Masson’s trichrome staining of a pancreas with low-to-moderate stromal content showing ADM and fibrosis (KC pig, left) and a highly stromal pancreas with extensive ADM and dysplastic changes (KPC pig, right). **(E–H)** Representative immunohistochemical staining of pancreatic lesions from KPC pigs showing stromal expression of vimentin (**E**) and αSMA (**F**), and epithelial expression of GATA6 **(G)** and E-cadherin (**H**). **(I)** KRT19 staining of a KPC pig pancreas demonstrating ADM, low-grade PanIN, fibrosis, and inflammatory infiltrates. **(J)** MIST1 staining of a KC pig pancreas indicating loss of acinar cell identity. **(K)** Relative mRNA expression of *KRAS4A*, *KRAS4B*, *TP53^Δ^*^152^*, circTP53* and *p16*, together with DNA methylation levels of the *p16* promoter in tumour tissue (T) compared with healthy pancreas (N).

### The porcine PDAC model shows consistent and early pancreatic tumour initiation and progression

The disease course was monitored for up to eight months in KPC pigs and one year in KC pigs. To study pancreatic tumour progression, KC pigs (n= 3 per time point) and KPC pigs (n= 4-5 per time point) were euthanised at 1–2 months, 4–6 months, 8 months, and, for KC pigs, also at one year. The severity and onset of symptoms varied among individual KPC pigs, with the first signs of disease, such as weight loss and apathy, occurring as early as 2-4 months in some individuals. In contrast, KC pigs showed no symptoms, even at one year of age. Necropsy results from control pigs (6 to 8 months old) expressing iCre only showed that expression of iCre alone did not cause any abnormalities. The choice of iCre driver-line did not significantly affect the timeline of PDAC progression. The latter was, however, strongly influenced by the presence or absence of mutant *TP53* as demonstrated by the significantly shorter lifespan of KPC pigs (up to a maximum of 8 months) compared to KC pigs and WT controls. At necropsy, macroscopic pancreatic tumours were already detectable in KPC pigs as early as one month of age (Figure 1B).

Histological analysis at all necropsy time points demonstrated that KC pigs exhibited only acinar-to-ductal metaplasia (ADM) (Figure 1C), with two exceptions (aged 1 and 5 months), in which individual tumours also displayed focal low-grade pancreatic intraepithelial neoplasia (PanIN). At one month of age most lesions showed multifocal ADM. As the pigs aged, the extent of ADM became more pronounced and widespread (Figure 1C), indicating a progressive expansion of these lesions over time. All lesions collected from KC pigs exhibited a firm, hardened texture on macroscopic examination.

In KPC pigs, tumour burden increased over time, with multiple pancreatic tumours reaching sizes of up to 4 cm (Figure 1B). Tumours from one to eight months old KPC pigs exhibited both extensive ADM and focal PanIN lesions (Figure 1C), with inflammatory reactions and periductal fibrosis becoming more prevalent from four months of age (Figure 1C). This pattern was observed both across different individuals and among distinct tumours within the same animal. In addition, a low-grade intraductal papillary mucinous neoplasm (IPMN) was detected in a single KPC pig at 4 months of age (Figure 1C).

### Immunohistological and transcriptomic analyses reveal tumour heterogeneity in porcine PDAC

Human PDAC is characterised by major transcriptional subtypes, including classical and mesenchymal/basal-like phenotypes and by a complex tumour microenvironment (TME) comprising stromal, immune, and extracellular matrix components^33–35^. The dense tumour stroma is one of the most prominent histological features of PDAC and contributes substantially to tumour architecture and biological heterogeneity. Classical epithelial features include glandular differentiation and expression of markers such as cytokeratin 19 (KRT19), E-cadherin and GATA6 (an endodermal lineage-specifying transcription factor) ^36^, whereas mesenchymal/basal-like features are associated with reduced epithelial differentiation and increased expression of markers linked to EMT and invasive behaviour, including vimentin and S100A2. In addition, stromal markers, including alpha-smooth muscle actin (αSMA), reflect the presence of activated fibroblasts and pancreatic stellate cells within the tumour microenvironment.

Cancer-associated fibroblasts (CAFs) are the main component of the desmoplastic reaction and play a critical role in the secretion of extracellular matrix components essential for forming a fibrotic and hypovascular TME. To assess stroma deposition in porcine pancreatic lesions, Masson’s trichrome staining was performed on tumour sections from KC and KPC pigs. Qualitative histological evaluation showed that KPC tumours consistently exhibited more prominent collagen-rich stromal areas than KC lesions in the examined samples. Representative examples are shown in Figure 1D. Although stromal deposition appeared qualitatively more prominent in KPC than in KC lesions, the expression of epithelial and stromal markers was highly heterogeneous among tumours isolated from KC and KPC pigs of different ages, as well as between distinct tumours from the same individual animal. This variability indicates pronounced intra- and inter-individual heterogeneity, closely resembling the molecular and histopathological diversity observed in human PDAC. Consistent with this, immunohistochemical analysis revealed a complex TME in porcine pancreatic tumours, characterised by the coexistence of epithelial tumour compartments and stromal components. Several tumours exhibited strong vimentin staining and/or stromal αSMA staining (Figure 1E,F) indicating the presence of mesenchymal/stromal components, including activated fibroblasts and pancreatic stellate cells. In parallel, tumour epithelial cells expressed classical PDAC markers, including GATA6 (Figure 1G), E-cadherin (Figure 1H), and KRT19 (Figure 1I) in progressive pancreatic lesions from both KC and KPC pigs.

Consistent with a ductal tumour phenotype, all pancreatic tumour sections showed strong KRT19 staining irrespective of genotype (Figure 1I). In lesions with extensive acinar-to-ductal metaplasia, immunohistological examination revealed loss of the acinar differentiation marker MIST1 (Figure 1J), indicating acinar cell reprogramming into ductal-like cells and progression toward a more advanced tumour state. The upregulation of *E-cadherin*, *GATA6*, and *αSMA* was also detected by qRT-PCR analysis of bulk tumour samples (Supplementary Figure 1A). However, because these analyses were performed on bulk tissue, increased αSMA expression most likely reflects the contribution of stromal cells rather than tumour epithelial-cell expression. Similarly, *vimentin* was upregulated in bulk samples (Supplementary Figure 1A), but immunohistochemistry indicated that this expression was at least partly attributable to stromal or CAF-associated compartments (Figure 1E). Interestingly, we observed a significant overexpression of *S100A2* mRNA in porcine tumour samples (Supplementary Figure 1A), a marker implicated in epithelial-mesenchymal transition (EMT), invasiveness, and metastatic potential in pancreatic cancer patients^37^. Furthermore, qPCR analysis showed increased expression of *CD44* isoforms in bulk tumour samples relative to normal pancreatic tissue, with the most pronounced upregulation observed for the standard isoform *CD44s*, whereas the analysed CD44 variant (*CD44v*) isoforms showed lower expression levels (Supplementary Figure 1A). While CD44s may originate from multiple cell types, including activated T cells and stromal cells within TME, the upregulation of CD44v is more indicative of cancer stem-like cells ^38, 39^. Cancer stem cells are defined by their capacity for self-renewal and differentiation, which are pivotal factors in promoting tumour growth, progression, and therapy resistance^40^.

In conclusion, these findings indicate that progressive pancreatic lesions in pigs share several molecular and histopathological features with human PDAC, including expression of epithelial PDAC-associated markers such as KRT19, GATA6, E-cadherin, together with prominent tumour-associated stromal components marked by collagen deposition, vimentin and stromal αSMA expression. In addition, transcriptional changes associated with EMT, tumour progression and cancer stem cell-like phenotypes further support the relevance of this model for studying PDAC heterogeneity.

### Porcine pancreatic tumours exhibit early alterations in tumour suppressor networks and *KRAS* isoform expression

We had previously identified an upregulation of the *TP53Δ152* isoform *and circTP53*, both of which are absent in the mouse, in our pig osteosarcoma model^6^. Thus, we investigated whether these *TP53* isoforms were also differentially expressed in both KC and KPC pigs with pancreatic lesions. In addition, we measured the expression of the p53 family members *TP63* and *TP73*, along with *p16*. These are key regulators of cell cycle control, differentiation and tumour suppression including pancreatic cancer^41, 42^. RNA analysis showed a significant upregulation of *p53Δ152, circTP53*, and *TP63ΔN* and a downregulation of *TP73* and *p16* in tumour samples (Figure 1K and Supplementary Figure 1B, C). The expression of *TP63ΔN* in some tumour samples suggests the presence of the basal-like/squamous PDAC subtype^43, 44^. To determine if the altered expression was due to epigenetic regulation, the promoter methylation of *TP53Δ152* and *p16* was analysed. A clear correlation between promoter methylation and mRNA expression was observed; lower methylation of *TP53Δ152* promoter resulted in increased *TP53Δ152* expression, while increased promoter methylation of the *p16* corresponded to reduced *p16* expression in tumour samples (Figure 1K).

In parallel, we investigated alterations in oncogenic *KRAS* signalling at the level of mRNA isoform expression. *KRAS* produces two splice variants, *KRAS4A* and *KRAS4B*, which differ in their C-terminal membrane-targeting sequences and intracellular localisation. In the context of the *KRAS^G12D^* mutation, both isoforms harbour the activating substitution. While *KRAS4B* is generally the predominant isoform, increased expression of *KRAS4A* has been reported in several cancers^45^. Relative to healthy pancreatic tissue, porcine pancreatic tumours exhibited significant upregulation of both *KRAS4A* and *KRAS4B* mRNA expression. In contrast, healthy pancreatic tissue showed only weak *KRAS4B* expression and an absence of detectable *KRAS4A* transcripts (Figure 1K).

Together, these findings demonstrate that porcine pancreatic lesions exhibit early and coordinated alterations in both tumour suppressor pathways and oncogenic KRAS signalling. Given that KC pigs predominantly developed acinar-to-ductal metaplasia (ADM), these molecular changes appear to arise at very early stages of pancreatic tumorigenesis, underscoring the utility of the porcine model for dissecting early events in PDAC development.

### Early invasive growth and lymphatic dissemination in porcine PDAC

Consistent with the aggressive biological behaviour of human PDAC, a majority of KPC pigs (n = 15) and a subset of KC pigs (n = 2) developed local lymph node (LN) metastases, as confirmed by KRT19-positive tumour cells in regional LNs (Figure 2A). Analysis of extracellular vesicle (EV)-enriched plasma fractions revealed increased levels of EV-associated mutant *KRAS* transcripts in pigs presenting LN metastases (Figure 2B), as quantified by qPCR. These findings indicate that the porcine PDAC model recapitulates the release of tumour-derived RNA via circulating EVs observed in human pancreatic cancer^46^ and supports its utility for longitudinal liquid biopsy monitoring. Notably, lymphatic dissemination was already detectable in two one-month-old KPC pigs, indicating that metastatic competence can arise at very early stages of tumour development. In one of these animals, clear evidence of vascular invasion was observed (Figure 2C), together with the presence of single invasive tumour cells infiltrating the surrounding tissue (Figure 2D).

**Figure 2.**
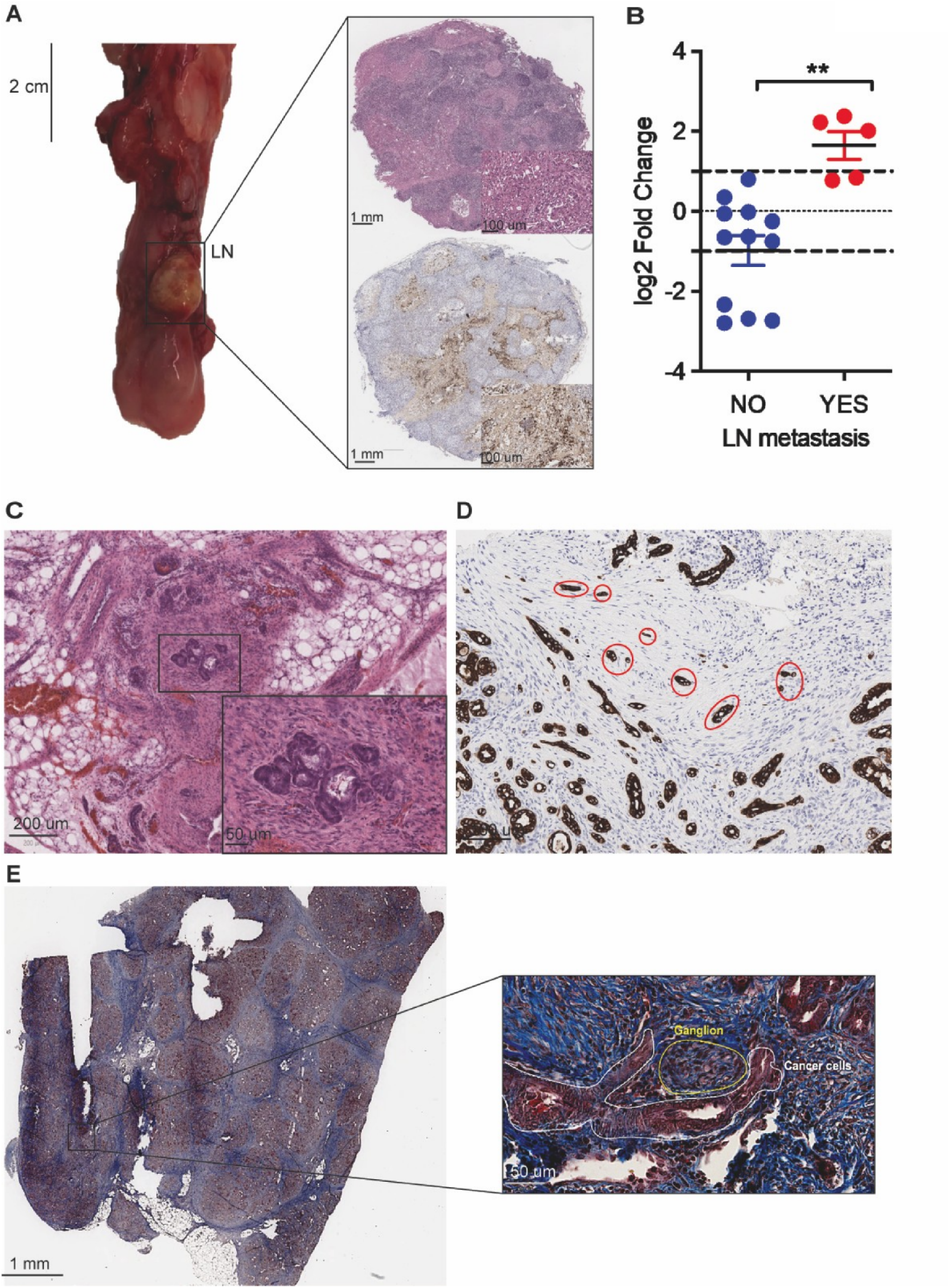
Tumour progression, invasion, metastasis, and circulating tumour-derived EV-associated RNA in a porcine PDAC model. **(A)** Macroscopic image of the pancreas from a KPC pig showing a primary pancreatic tumour and an adjacent lymph node (left), and representative H&E staining and KRT19 immunohistochemistry of the lymph node demonstrating KRT19-positive metastatic tumour cells (right). **(B)** Analysis of extracellular vesicle (EV)-enriched plasma-derived cfRNA demonstrating increased levels of circulating mutant *KRAS* transcripts in pigs with lymph node metastases, indicating release of tumour-derived EV-associated RNA and supporting its utility as a liquid biopsy biomarker in porcine PDAC. **(C)** Representative H&E-stained section showing vascular invasion by tumour cells in a KPC pig pancreas. **(D)** KRT19 immunohistochemical staining revealing single invasive tumour cells infiltrating surrounding tissue. **(E)** Masson’s trichrome staining of a KPC pig pancreas illustrating perineural invasion (PNI), characterised by tumour cells encasing a nerve ganglion.

### Early perineural invasion in the porcine PDAC model

Perineural invasion (PNI), a hallmark feature of early human PDAC but typically absent in the KPC mouse model^47^, was also detected in the porcine model. Histological examination revealed tumour cells surrounding nerve ganglia in KPC pigs as early as one month of age (Figure 2E). Although PNI was not as extensive as commonly observed in advanced human PDAC, its early presence further supports the translational relevance of the porcine model.

### Early porcine PDAC exhibits an organised and functionally active T-cell immune landscape

To characterise the immune landscape of porcine PDAC at both the organisational and functional levels, single-cell RNA sequencing (scRNA-seq) was performed on tumour-adjacent lymph nodes and flow cytometry on primary tumours. scRNA-seq analysis of tumour-draining lymph nodes from a four-month-old KPC pig identified 13 distinct immune cell clusters (Figure 3A). These included four B-cell populations expressing CD19, CD21, and CD79B, corresponding to resting, activated, cycling B cells, and plasma cells (Figure 3B, C). In addition, an NK cell cluster expressing KLRB1 and KLRK1 (encoding CD161 and NKG2D, respectively) and a myeloid cell cluster comprising dendritic cells and macrophages were detected. T cells expressing CD3E, CD3D, and CD2 segregated into seven subclusters, including two CD4+CD8A− T-cell populations and one CD4+CD25+Foxp3+ regulatory T-cell cluster (Figure 3C, D). Notably, the regulatory T-cell cluster contained a subset of cells expressing interferon-γ (IFN-γ) and tumour necrosis factor (TNF), indicating functional heterogeneity within this compartment. In addition, perforin-expressing CD8B+ T cells were distributed across two cytotoxic CD8+ T-cell clusters. These findings are consistent with previous reports describing diverse T-cell subsets with distinct activation states in human PDAC-draining lymph nodes ^48, 49^.

**Figure 3.**
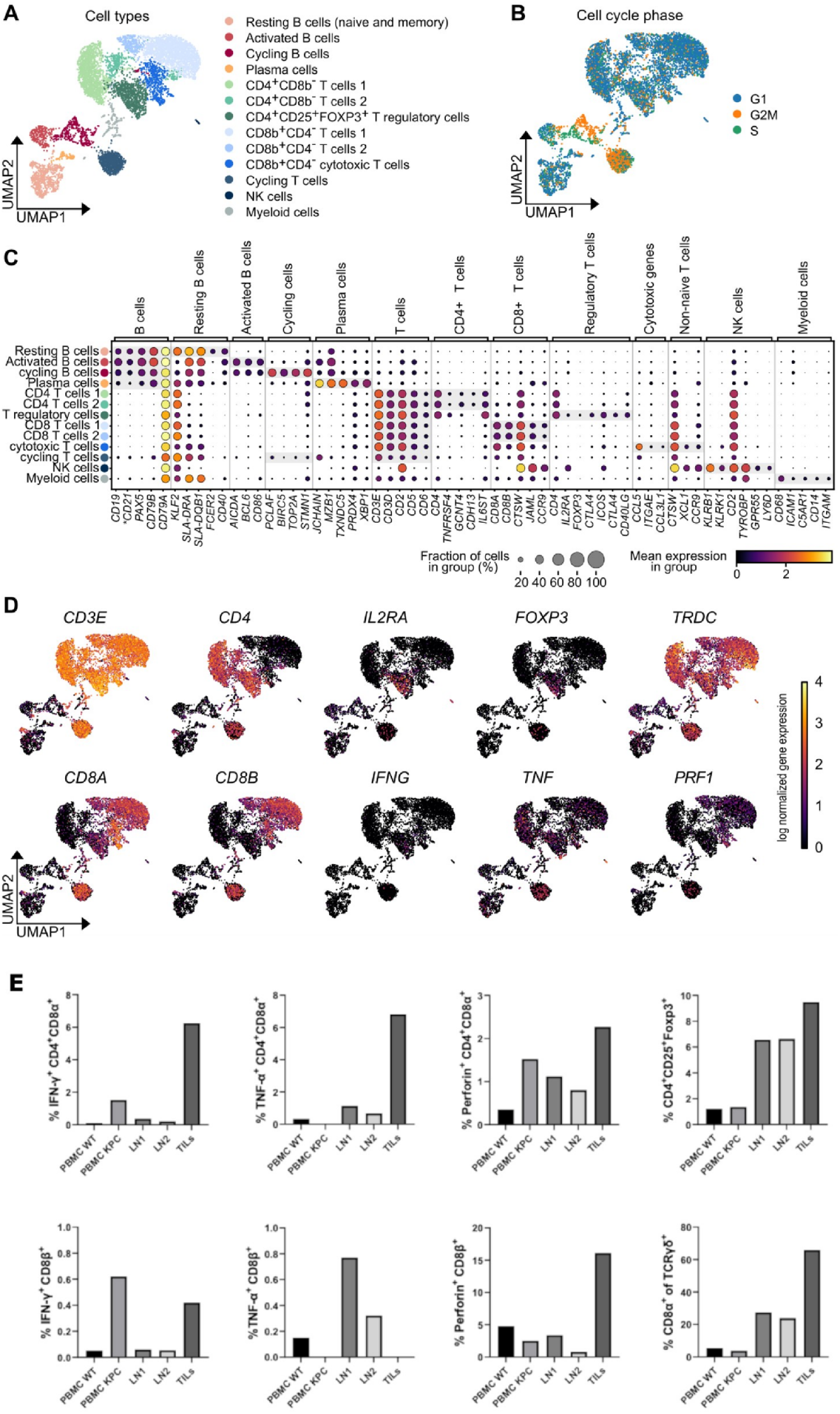
Organisational and functional immune landscape of porcine PDAC. Single-cell RNA sequencing (scRNA-seq) of tumour-draining lymph nodes and flow cytometric analysis of primary tumours from a four-month-old KPC pig. **(A)** UMAP representation of immune cell populations identified by scRNA-seq. **(B)** UMAP showing cell-cycle phase distribution across immune cell clusters. **(C)** Dot plot of selected marker genes used for immune cell type annotation, including B-cell, myeloid, NK-cell, and T-cell populations. **(D)** UMAP projection of T-cell subset markers, including CD4⁺CD8A⁻ T cells, CD4⁺CD25⁺FOXP3⁺ regulatory T cells, CD8⁺ cytotoxic T cells, γδ T cells, and perforin-expressing T-cell populations. **(E)** Flow cytometric analysis of tumour tissue, tumour-adjacent lymph nodes, and peripheral blood showing IFN-γ-, TNF-, and perforin-producing CD4⁺CD8α⁺ and CD8β⁺ T cells, demonstrating functional effector and cytotoxic T-cell activity within the tumour microenvironment.

To assess whether these immune populations infiltrate primary tumours and display functional activity, flow cytometry was performed on tumour tissue, tumour-adjacent lymph nodes, and peripheral blood from four-month-old KPC pigs and wild-type littermate controls.

Tumour tissue was infiltrated by cytotoxic CD8β+ and effector CD4+CD8α+ T cells, along with CD4+CD25+Foxp3+ regulatory T cells. Within the tumour microenvironment, CD4+CD8α+ T cells producing IFN-γ (6.25%) and TNF (6.82%) were detected, together with perforin-producing CD8β+ T cells (Figure 3E).

The presence of functionally active effector and cytotoxic T cells suggests that early porcine PDAC lesions, predominantly characterised by ADM with few PanINs, exhibit features of an immunologically “hot” tumour phenotype. This interpretation is further supported by the detection of tertiary lymphoid structures (TLSs) in tumours from KPC pigs (Supplementary Figure S2), which may facilitate local immune activation and recruitment of effector T cells.

In contrast, clinically manifest human PDAC is generally considered immunologically “cold” ^50^, whereas the immune landscape of early-stage precursor lesions remains less well defined due to limited access to early human samples, despite increasing recent efforts to characterise these settings^51^.

### Establishment and molecular characterisation of porcine pancreatic cancer cell lines

Cancer cell lines are important tools to assess drug targets prior to *in vivo* interventions. Thus, we established cell lines from early KC and KPC pancreatic tumours. Given the stromal-rich nature of tumour samples from KPC pigs, we examined the presence of CAFs at early passage (P3) by a) performing PCR to detect the Cre-induced activation of the *KRAS^G12D^*mutant allele, which is indicative of cancer cells and does not occur in CAFs, b) by isolating tumour cells from one KPC/R26^mT/mG^ pig that also carried a Cre-activatable dual-fluorescent reporter gene^52^. Cells with expression of mutant *KRAS*/*TP53* should be green fluorescent, while all other cells would be red as shown in Supplementary Figure S3A.

In total, 17 cell isolates (nine from KPC and eight from KC pigs) were successfully cultured beyond passage 30 (hereafter referred to as porcine pancreatic cancer cell (PoPaCa) lines). PCR analysis revealed the presence of the unrecombined *KRAS* allele, indicative of CAFs (marked as LSL, in 8 of 17 PoPaCa lines). The remaining nine PoPaCa lines were deemed as CAF-free. PCR results also indicated an overrepresentation of the mutant KRAS allele in some cell lines or even a loss of the *KRAS^WT^* allele (PoPaCa-3 and -6) (Figure 4A).

**Figure 4.**
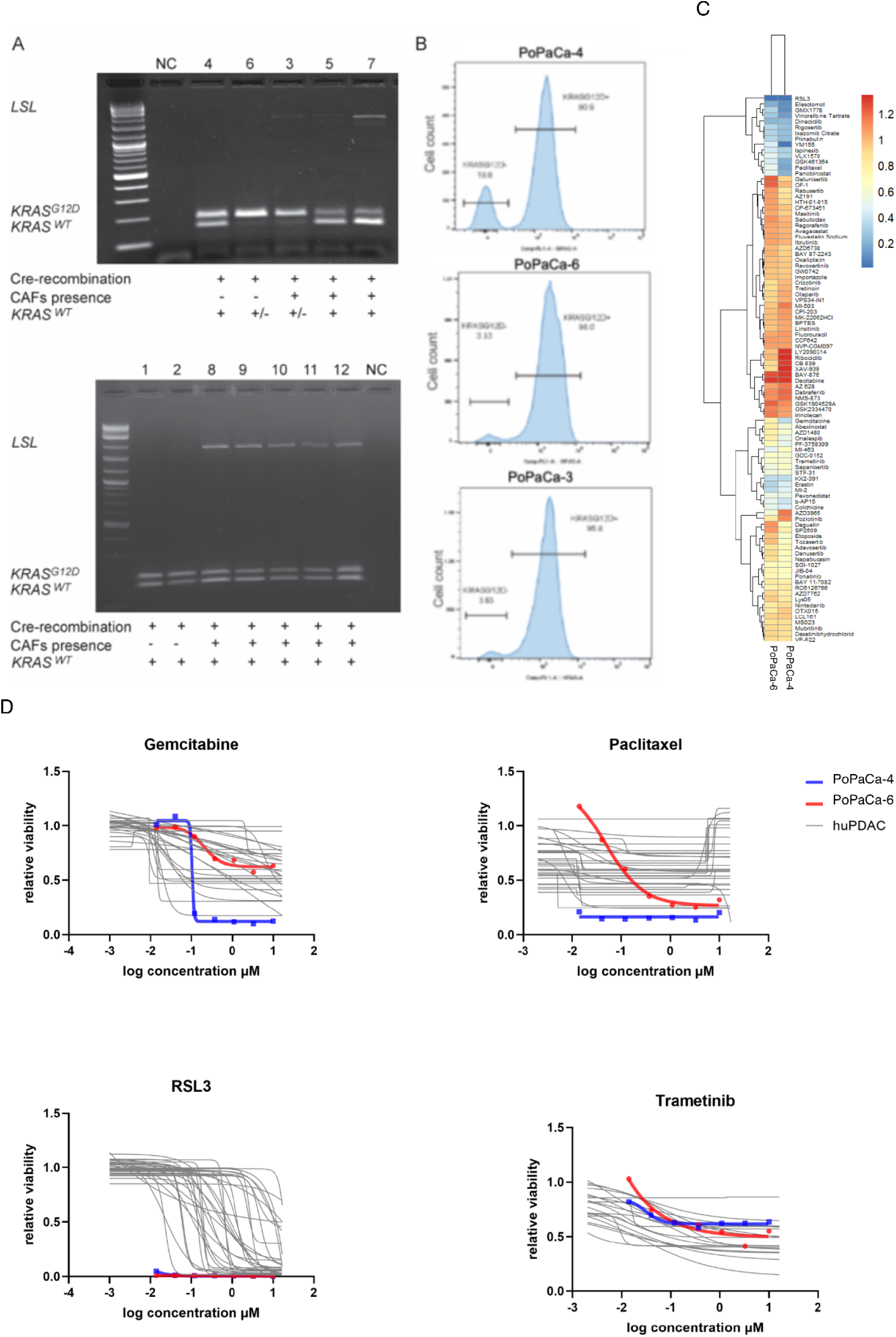
Genetic characterisation and drug response profiling of porcine pancreatic cancer cell lines. **(A)** PCR analysis of KRAS recombination in pancreatic cancer cell lines (PoPaCa) derived from independent KC and KPC tumours. Presence of the non-recombined *KRAS^LSL^* allele indicates contamination with cancer-associated fibroblasts (CAFs), whereas absence of the wild-type *KRAS* allele indicates loss of *KRAS^WT^* in tumour cells. **(B)** Flow cytometric quantification of *KRAS^G12D^*-positive and *KRAS^G12D^*-negative cells in PoPaCa cultures, demonstrating variable tumour cell purity and KRAS allelic composition among individual lines. **(C)** Clustered heatmap of area-under-the-curve (AUC) values from compound screening in PoPaCa-6 and PoPaCa-4. **(D)** Qualitative comparison of dose-response profiles of two CAF-free porcine PoPaCa lines and human PDAC cell lines. Relative viability curves are shown for PoPaCa-4 and PoPaCa-6, which differ in *KRAS* allelic status, together with publicly available human PDAC cell-line dose-response data from the CTD² dataset for gemcitabine, paclitaxel, RSL3 and trametinib.

Loss of *KRAS^WT^* and increased dosage of oncogenic *KRAS^G12D^* are frequently observed in human cancers and are associated with tumour progression^53, 54^. To determine whether similar allelic imbalances occur in porcine pancreatic tumours, we quantified *KRAS* allele frequencies in PoPaCa lines and primary tumour material. Consistent with the PCR data, loss of *KRAS^WT^*was confirmed in PoPaCa-3 and -6. At the protein level, analysis using a *KRAS^G12D^*-specific antibody demonstrated that *KRAS^G12D^* constituted approximately 80% of total KRAS protein in PoPaCa-4 (harbouring both alleles), and 96% in PoPaCa-3 and -6, confirming near-complete loss of *KRAS^WT^* in these lines (Figure 4B).

To assess whether allelic imbalance was detected already *in vivo* and not acquired during *in vitr*o culture, *KRAS* allele frequencies were analysed in bulk tumour samples and microdissected tumour epithelium. Four of fourteen microdissected tumours exhibited mutant *KRAS* allele frequencies approaching 100% (Supplementary Figure S3B). In corresponding bulk tumour samples, the relative abundance of mutant KRAS was lower (∼70%), consistent with dilution by stromal cells (Supplementary Figure S3B). Together, these data indicate that increased dosage of oncogenic *KRAS* can arise at early stages of porcine pancreatic tumour development.

To further characterise the molecular features of PoPaCa lines, bulk RNA sequencing was performed on twelve PoPaCa cell lines. Comparative transcriptomic analyses identified distinct gene expression signatures associated with tumour-intrinsic properties and CAF content, with pathway enrichment analyses indicating involvement of cancer-relevant metabolic and extracellular matrix–associated processes (Supplementary Figure S3C).

Collectively, these data indicate that porcine PoPaCa lines retain key genetic, transcriptional, and microenvironment-associated features of pancreatic tumour development, supporting their use as a translational platform for studying PDAC biology and for preclinical evaluation of targeted therapies.

### Drug response profiling of porcine pancreatic cancer cell lines identifies heterogeneous treatment responses

To explore the suitability of porcine pancreatic cancer cell lines for preclinical drug-response studies, we performed an exploratory high-throughput drug screen using two CAF-free PoPaCa lines. These lines differed in *KRAS* allelic status: PoPaCa-6 showed increased mutant *KRAS* expression and loss of the *KRAS^WT^* allele, whereas PoPaCa-4 retained *KRAS^WT^*. Given the limited number of cell lines analysed, these experiments were not intended to establish definitive genotype–response associations, but rather to assess whether porcine PDAC-derived cell lines show measurable and heterogeneous responses to clinically relevant and experimental compounds. The two PoPaCa lines were screened against a panel of 95 compounds targeting diverse pathways implicated in cancer development and progression (Supplementary Table S3). Of these compounds, 26% are approved for the treatment of pancreatic cancer or other malignancies, 22% are currently in Phase I–III clinical trials, and 52% are in preclinical development. Drug responses were summarised using area-under-the-curve (AUC) values (Figure 4C). Because AUC values depend on the tested concentration range and potency of each compound, they were interpreted primarily within individual drugs and between the two porcine cell lines, rather than as absolute measures of sensitivity or resistance across drug classes.

Overall, the drug-response profiles of PoPaCa-4 and PoPaCa-6 were broadly comparable, with most compounds eliciting similar responses in both cell lines. Both lines showed reduced viability in response to several compounds targeting cell-cycle regulation, cytoskeletal dynamics and epigenetic regulators. Responses were also observed for selected compounds targeting metabolic dependencies, such as GMX1778, an inhibitor of nicotinamide phosphoribosyltransferase (NAMPT). Conversely, several compounds within signalling-related categories, including selected PI3K/AKT/mTOR, MAPK and DNA-damage response inhibitors, showed limited activity under the tested conditions. These observations should be interpreted cautiously, as formal classification of sensitivity or resistance would require comparison with larger panels of porcine and human PDAC cell lines tested under matched experimental conditions.

Despite the overall similarity between the two lines, differential responses were observed for a subset of compounds. PoPaCa-4 showed stronger responses than PoPaCa-6 to several cytotoxic agents, including gemcitabine, paclitaxel and etoposide; inhibitors of cell-cycle or mitotic progression, including tozasertib, GSK461364 and rabusertib; the epigenetic regulator panobinostat; and metabolism-targeting compounds, including deguelin and GMX1778 (Figure 4C). Although these differences may reflect biological heterogeneity between the two porcine PDAC cell lines, the small number of lines analysed precludes direct attribution of these responses to *KRAS* allelic status alone.

To place the porcine drug-response profiles into the context of human PDAC, selected dose-response curves from the two porcine PDAC cell lines were plotted together with publicly available human PDAC cell-line data from the CTD² dataset^28, 29^. Relative viability of PoPaCa-6 and PoPaCa-4 is shown together with human PDAC cell-line dose-response curves for gemcitabine, paclitaxel, RSL3 and trametinib (Figure 4D). Because concentration ranges and screening conditions differed between datasets, this comparison was restricted to qualitative visual assessment and was not used for direct quantitative AUC-based analysis.

For gemcitabine, the response profiles of the porcine PDAC cell lines fell within the range observed for human PDAC cell lines, although the two porcine lines differed markedly from each other, with PoPaCa-4 showing a stronger reduction in viability than PoPaCa-6. A similar pattern was observed for paclitaxel, where PoPaCa-4 displayed a more pronounced dose-dependent decrease in viability, consistent with its lower AUC value in the porcine drug screen. Together, these findings indicate that PoPaCa-4 was more responsive than PoPaCa-6 to selected cytotoxic agents, including gemcitabine and paclitaxel, while both porcine cell lines remained within the qualitative response spectrum observed among human PDAC cell lines (Figure 4D).

A distinct response pattern was observed for the ferroptosis inducer and GPX4 inhibitor RSL3. Both porcine PDAC cell lines showed near-complete loss of viability already at the lowest tested concentration, whereas human PDAC cell lines displayed a broader range of dose-response profiles and generally required higher concentrations to reach comparable effects (Figure 4D). This observation requires validation in additional porcine PDAC cell lines and under matched screening conditions but suggests that ferroptosis induction may represent a potential vulnerability in these models.

In contrast, the porcine PDAC cell lines showed only partial responses to the MEK inhibitor trametinib. In qualitative comparison with human PDAC cell-line data, their dose-response curves resembled those of the less responsive human PDAC cell lines in the comparative dataset (Figure 4D). Further studies will be required to determine whether this reflects species-specific differences, pathway adaptation, *KRAS*-related signalling states or other molecular features of the individual porcine cell lines.

Together, these exploratory drug-response data indicate that porcine PDAC cell lines exhibit measurable and heterogeneous responses to clinically relevant and experimental compounds. The responses to selected cytotoxic agents, including gemcitabine and paclitaxel, qualitatively overlapped with the range observed in human PDAC cell lines, while the differential responsiveness of PoPaCa-4 and PoPaCa-6 highlights inter-line heterogeneity. The pronounced response to RSL3 and partial response to trametinib identify testable drug-response patterns for future validation in larger porcine PDAC cell-line panels.

## Discussion

Pancreatic cancer is a devastating disease, often presenting at an advanced stage and so far with limited effective treatment options, resulting in poor patient outcomes. Recent advances in personalised RNA-based cancer vaccines^4, 55^ show promising results, but open questions remain here, e.g. why some patients respond, and others do not. While therapeutic interventions might be most effective at early stages of the disease, current diagnostic methods remain suboptimal despite recent progress, such as the PAC-MANN assay, which improves detection of stage I PDAC^56^.The new translational PDAC model, presented here, might help fill these gaps. Pigs have emerged as a valuable option due to their similarities with humans in body size, anatomy, physiology and pathophysiological responses^57^. They are widely used models for toxicity studies, medical device testing, imaging, and surgical interventions. Their value as preclinical and translational research tools have previously been reported ^6–11, 13, 58–60^.

Although pigs carrying inducible *KRAS^G12D^* and *TP53^R^*^167^*^H^* mutations^30, 31^, have been available for some time, due to the lack of pancreas-specific iCre-driver lines, attempts to create a porcine PDAC model had proven technically difficult^15, 16^. Local induction of pancreatic tumours by injecting adenoviral vectors expressing Cre recombinase into the pancreas of Oncopigs has been reported^17, 61^. However, the utility of the model is limited due to rapid disease progression (< 2 weeks), lack of precursor lesions and inflammatory reaction. Recently, we reported the successful development of two *PTF1A^Cre^* driver lines (*PTF1A^5’iCre^*and *PTF1A^3’iCre^*)^18^. The transcription factor Ptf1a is essential for maintaining mature acinar identity, and its loss promotes tumorigenesis^62^. In contrast to iCre insertion at 3’UTR, placing iCre at the 5’ end of the *PTF1A* gene resulted in a heterozygous knockout, possibly affecting tumour development. However, no correlation in the severity of pancreatic lesions was observed when comparing the two *PTF1A*-iCre pig lines.

Both iCre-lines successfully activated mutant *KRAS^G12D^*and *TP53^R^*^167^*^H^*in a tissue-specific manner resulting in a reproducible porcine PDAC model that recapitulates several clinically relevant features of human PDAC and shows distinct progression kinetics in KC and KPC animals. KC pigs showed fewer tumour lesions (mainly ADMs) and a more extended latency period. Concomitant expression of *KRAS^G12D^* and *TP53^R^*^167^*^H^* leads to disease progression from preinvasive to invasive and metastatic PDAC. Macroscopically visible nodular lesions were observed in all KPC piglets, with histological features consistent with human acinar-ductal metaplasia (ADM), pancreatic intraepithelial neoplasia (PanINs) at one month of age and invasive PDAC with metastasis to lymph nodes. The early metastasis to regional lymph nodes mirrors the aggressive nature of human PDAC, which is the leading cause of cancer-related death. Yet our understanding of the metastatic process remains limited due to its multi-step nature and the influence of the *in vivo* microenvironment^63^. The ability of the PDAC pig model to recapitulate early metastatic events provides a robust platform to study the initial stages of cancer cell dissemination and to identify key drivers of metastasis. In addition, recent imaging studies in this model have revealed stages of PDAC progression not typically observed in humans, providing a unique opportunity to study tumour progression in a controlled setting and develop more effective early diagnostic and therapeutic strategies.

To aid interventional studies, the model was established in a porcine hybrid line, with adults reaching 40 to 50 kg at 12 months of age. This reduces costs and facilitates intervention studies with drug dosages like those administered to humans while still being large enough for testing human-scale devices.

PDAC is a heterogeneous disease with diverse molecular subtypes characterised by a highly complex and diverse tumour microenvironment (TME) composed of stroma, vascular, immune and extracellular matrix components^34^. Understanding this complexity is essential for the development of effective therapeutic strategies. Recent studies have highlighted the critical role of oncogenic KRAS dosage and signalling in shaping cell morphology, plasticity, histopathology, and disease progression. In particular, mesenchymal PDAC subtypes exhibit increased *KRAS^G12D^*amplification and expression, which correlate with enhanced tumour aggressiveness, metastatic potential, and resistance to standard chemotherapy, underscoring a major unmet clinical need^53, 64^. The porcine PDAC model presented here offers a valuable platform to study these molecular subtypes and their associated TME. Molecular analyses further support its translational relevance, as porcine tumours recapitulate key features of human PDAC, including epigenetic silencing of *p16*, increased mutant KRAS copy number, and altered expression of *KRAS* and *TP53* isoforms. Consistent with human data, these alterations appear to occur early during tumour development^65^. Notably, overexpression of *ΔNp63*, a key driver of basal-like/squamous PDAC subtype, was observed in porcine tumours. This isoform, together with *TP53Δ152* and *circTP53*, is absent in murine models^43^, highlighting the added value of the porcine system. Collectively, these findings demonstrate the ability of the model to capture early molecular events in PDAC initiation and progression, thereby facilitating studies on early detection and therapeutic interventions. In addition, extracellular vesicle (EV)-associated tumour-derived RNA represents a promising liquid biopsy biomarker in pancreatic cancer^66^. Detection of mutant *KRAS* transcripts in EV-enriched plasma from PDAC pigs enables longitudinal monitoring of tumour-derived biomarkers in a clinically relevant large animal model.

Transcriptomic analysis of PoPaCa lines from KC and KPC tumours revealed gene expression profiles aligned with pancreatic cancer-associated pathways. Differentially expressed genes indicated changes in lipid metabolism, cell signalling, and tumour-specific mechanisms, reinforcing the model’s relevance to human disease. Importantly, PoPaCa lines containing cancer-associated fibroblasts (CAFs) exhibited gene upregulation associated with extracellular matrix modulation and tumour-stroma interactions. This aligns with human PDAC, where CAFs contribute to increased collagen formation, extracellular matrix stiffness, and enhanced tumour cell migration.

To further enhance the translational value of the model, crossing KPC and KC pigs with dual-fluorescent reporter pigs^52^ will enable precise isolation and separation of CAFs and tumour cells. This strategy will facilitate the study of their individual contributions to tumour biology and drug responsiveness *in vitro*, while *in vivo*, GFP expressing tumour cells will enable tracking of metastatic dissemination and provide insights into the dynamics of PDAC progression.

The immunosuppressive TME is a unifying feature of all PDAC subtypes, which hinders the effectiveness of immunotherapies^67^. The presence of regulatory T cells within the tumour of the KPC pigs indicates immunosuppression, which can potentially compromise the anti-tumour activities of the effector CD4+ and CD8+ T cells. Similarly, in the Oncopig model^61^, adenoviral Cre-induced tumours exhibit a strong cytotoxic immune response, dominated by CD8+ T cells. However, this response was partially counterbalanced by increased Foxp3+ regulatory T cells and immunosuppressive molecules, such as IDO1, CTLA4, and PDL1. These findings highlight the importance of studying immune suppression in PDAC. As the porcine immunome and inflammasome are similar to those of humans ^57^, the PDAC pig model should be able to support the preclinical evaluation of immunotherapies aimed at restoring *in vivo* tumour-directed cytotoxicity.

The exploratory drug-response analysis provides initial evidence that porcine PDAC-derived cell lines may be useful for modelling pharmacological heterogeneity observed in human pancreatic cancer. Human PDAC is characterised by substantial intertumoural and intratumoural heterogeneity, which contributes to variable responses to cytotoxic chemotherapy and targeted therapies^33, 35, 68^. Consistent with this, the two CAF-free PoPaCa lines analysed here displayed distinct response profiles across the compound panel, despite both originating from genetically engineered *KRAS*-driven pancreatic lesions. Although the limited number of porcine cell lines precludes definitive conclusions regarding sensitivity or resistance patterns, these findings indicate that PoPaCa lines capture measurable and heterogeneous drug responses that are relevant for future preclinical testing.

Qualitative comparison with publicly available human PDAC cell-line data from the CTD² dataset^28, 29^ further supported this interpretation. The responses of the porcine PDAC cell lines to the standard chemotherapeutic agents gemcitabine and paclitaxel broadly reflected the range observed for human PDAC cell lines, although clear differences were observed between the two porcine lines. This is consistent with the well-established variability in chemotherapy response among human PDAC models and patients. In mouse KPC-derived systems, variable responses to gemcitabine-based treatment have also been reported, reflecting the strong influence of tumour genotype, differentiation state and microenvironmental context on therapeutic outcome. Thus, the heterogeneous responses observed in PoPaCa lines are compatible with the broader biology of PDAC rather than indicating a uniform drug-response phenotype.

One of the most notable observations was the pronounced sensitivity of both porcine PDAC cell lines to the ferroptosis inducer and GPX4 inhibitor RSL3. Ferroptosis has emerged as a potential therapeutic vulnerability in pancreatic cancer, particularly in relation to altered redox metabolism and lipid peroxide detoxification pathway^69^. For example, inhibition of GOT1, a key component of the non-canonical glutamine metabolism pathway used by PDAC cells to maintain redox balance, has been shown to promote ferroptotic cell death in pancreatic cancer models. In addition, therapy-resistant and mesenchymal-like cancer cell states have been associated with increased dependency on GPX4-mediated lipid peroxide detoxification, providing a mechanistic rationale for sensitivity to GPX4 inhibition^70, 71^. The strong response to RSL3 in the two PoPaCa lines is therefore biologically plausible and may reflect a shared ferroptosis-prone cell state, potentially linked to mesenchymal-like or therapy-resistant features, rather than differences in *KRAS* allelic status alone. However, because only two porcine lines were tested and the comparison with human data was qualitative, this finding should be considered hypothesis-generating and will require validation in larger porcine PDAC cell-line panels, molecular profiling of ferroptosis-related pathways, and ultimately *in vivo* studies.

In contrast, the porcine PDAC cell lines showed only partial responses to the MEK inhibitor trametinib, with dose-response curves resembling those of the less responsive human PDAC cell lines in the comparative dataset. This observation is in line with the known limited efficacy of single-agent MAPK pathway inhibition in PDAC. Although oncogenic *KRAS* activates RAF–MEK–ERK signalling, both human and mouse studies have demonstrated that MAPK pathway blockade can be limited by adaptive pathway rewiring, compensatory signalling and tumour-state-dependent resistance mechanisms, including AP1/FRA1-associated modulation of responsiveness to RAF–MEK–ERK cascade inhibition^25, 72, 73^. The limited response to trametinib observed in PoPaCa lines is therefore consistent with the broader challenge of targeting *KRAS* downstream signalling in PDAC, but further studies are needed to define the molecular basis of this phenotype in the porcine model.

The two PoPaCa lines also differed in *KRAS* allelic status, with one line showing loss of the *KRAS^WT^* allele and increased mutant *KRAS* expression, whereas the other retained *KRAS^WT^*. In human PDAC, *KRAS* dosage, allelic imbalance and mutant *KRAS* amplification have been linked to tumour evolution, transcriptional state, metabolic adaptation and therapeutic vulnerabilities^53^. However, the present study was not designed to establish a causal relationship between KRAS allelic status and drug response. Therefore, the observed differences between the two PoPaCa lines should be interpreted as evidence of biological heterogeneity rather than as proof of *KRAS*-dependent therapeutic response patterns.

Together, these findings suggest that porcine PDAC-derived cell lines can reproduce selected aspects of the pharmacological diversity observed in human pancreatic cancer models. Their qualitative overlap with human PDAC cell-line responses to standard chemotherapeutic agents, combined with the pronounced sensitivity to ferroptosis induction and limited response to MEK inhibition, identifies testable therapeutic hypotheses for future studies. Larger panels of porcine PDAC cell lines, integrated with genomic and transcriptomic profiling and matched cross-species drug-response analyses, will be required to determine whether specific molecular states or genetic alterations predict therapeutic sensitivity in this model.

## Conclusion

The development of a genetically engineered porcine model of pancreatic tumour development represents an important step toward modelling PDAC in a clinically relevant large animal system. Pancreas-specific activation of oncogenic *KRAS^G12D^*, alone or in combination with mutant *TP53R*^167^*^H^*, induced progressive pancreatic lesions that recapitulate several key features of human PDAC, including ADM, PanIN formation, invasive growth, perineural invasion, regional lymph node dissemination, stromal remodelling and tumour heterogeneity. Molecular and immunological analyses further demonstrated alterations in KRAS signalling, tumour suppressor networks, stromal composition and immune-cell organisation that are relevant to human pancreatic cancer biology.

The establishment of porcine pancreatic cancer cell lines and their exploratory drug profiling provide additional tools for studying tumour-intrinsic heterogeneity and therapeutic response. Together with the detection of tumour-associated circulating *KRAS* transcripts and the potential for longitudinal imaging and tissue sampling, this model offers a platform for investigating early tumour evolution, biomarker development and preclinical therapeutic strategies beyond the limitations of rodent models.

## Acknowledgments

The authors would like to thank Sara Robin and Benjamin Strickland for their technical support and Steffen and Viola Löbnitz, Gerhard Kammermeier, Andreas Sohn and Konrad Praller for animal husbandry. We would also like to acknowledge the Center for Advanced Light Microscopy (CALM, Technical University of Munich, Germany) for providing confocal laser scanning microscopes.

## Data availability statement

The original contributions presented in the study are included in the article and supplementary material. Further inquiries can be directed to the corresponding author Dr. Tatiana Flisikowska

