## Supplemental Data 1 for "A genetically engineered porcine model of pancreatic ductal adenocarcinoma captures the biological complexity of the human disease": PDAC Supplementary information.pdf

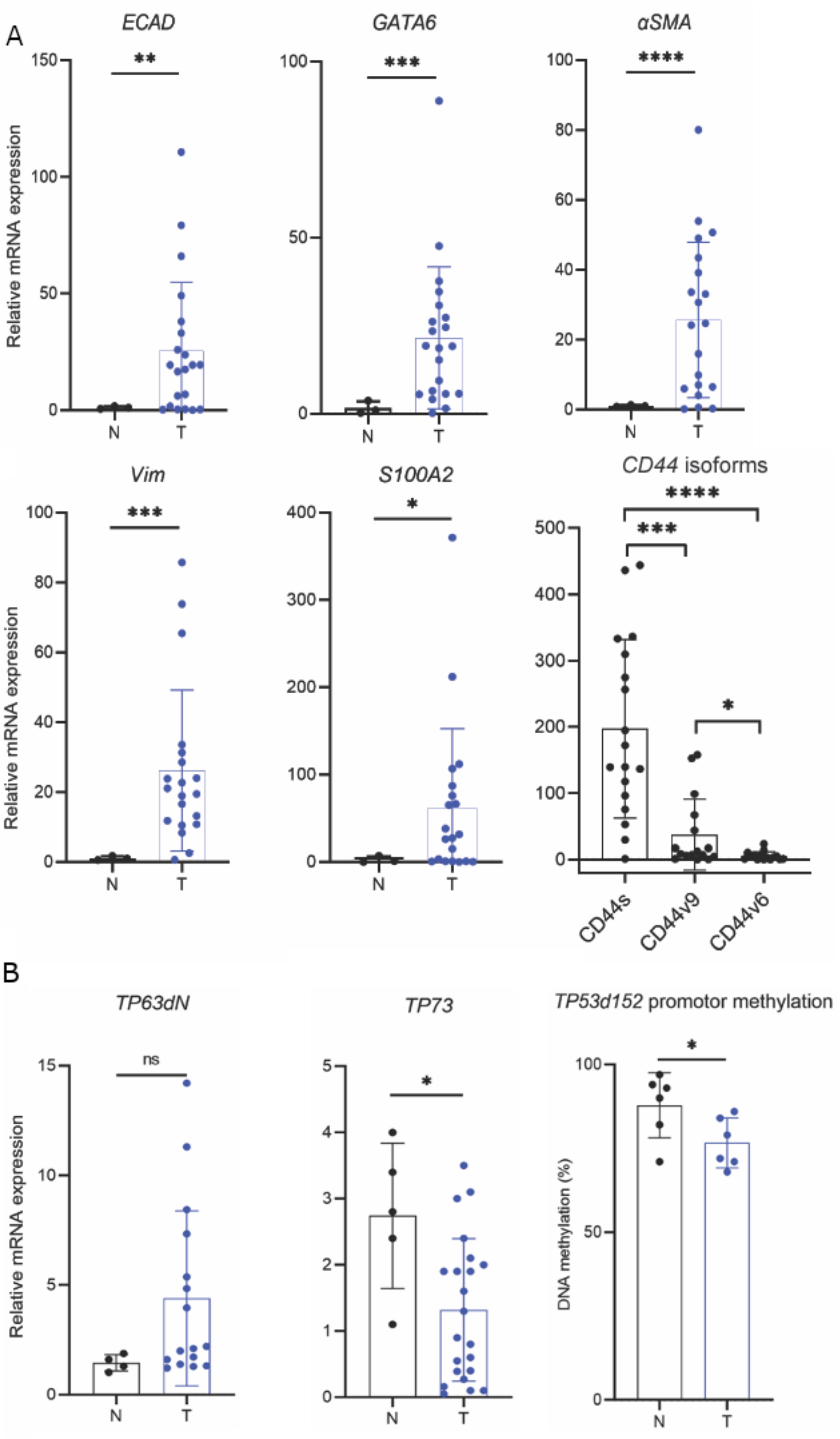


**Supplementary Figure S1 | Gene expression and epigenetic alterations in porcine PDAC tissues.**

**(A)** Relative mRNA expression levels of epithelial (*ECAD*, *GATA6*) and mesenchymal/stromal markers (*αSMA*, *VIM*, *S100A2*) in normal pancreas (N) and tumour tissue (T), determined by RT-qPCR. CD44 isoforms showed differential regulation, with predominant upregulation of the standard isoform CD44s in tumour samples. Expression of each isoform was normalised to the corresponding isoform expression level in normal pancreatic tissue, which served as the calibrator.

**(B)** Analysis of *TP63ΔN* and *TP73* expression indicates variable activation of TP53 family members, while *TP53Δ152* promoter methylation is reduced in tumours compared to normal tissue, consistent with increased expression.

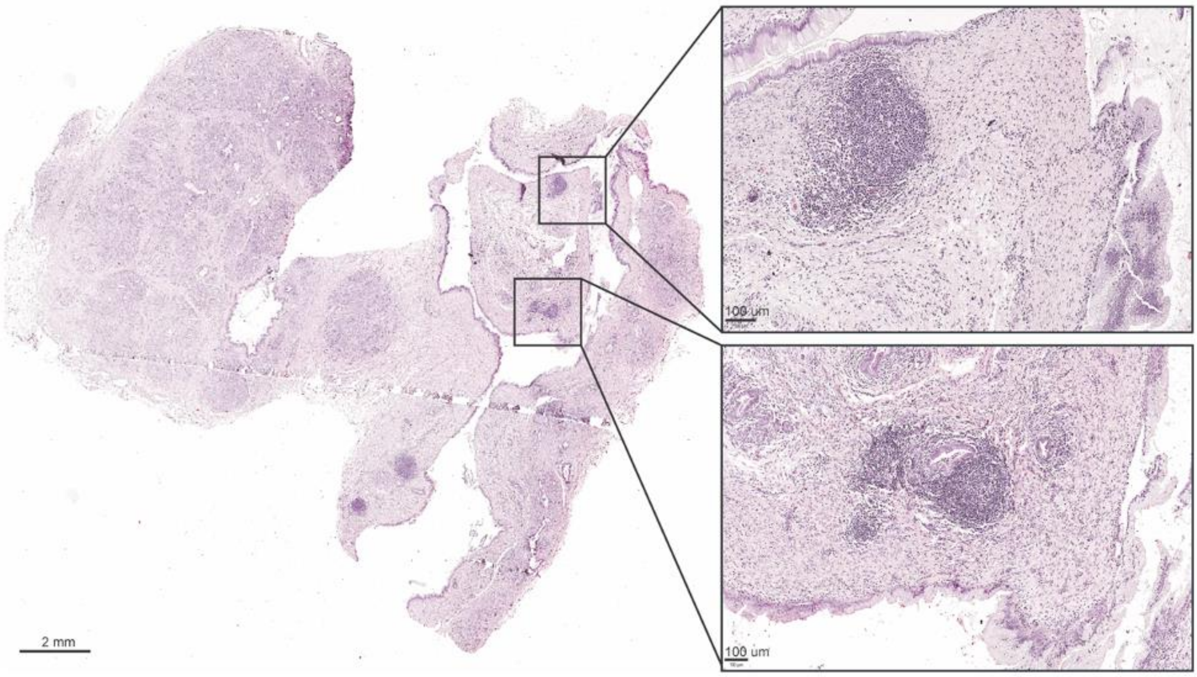


**Supplementary Figure S2 | Tertiary lymphoid structures in porcine PDAC.**
Haematoxylin and eosin (H&E) staining of a pancreatic tumour from a KPC pig showing dense fibrotic stroma with clearly identifiable tertiary lymphoid structures (TLS) within the tumour microenvironment.

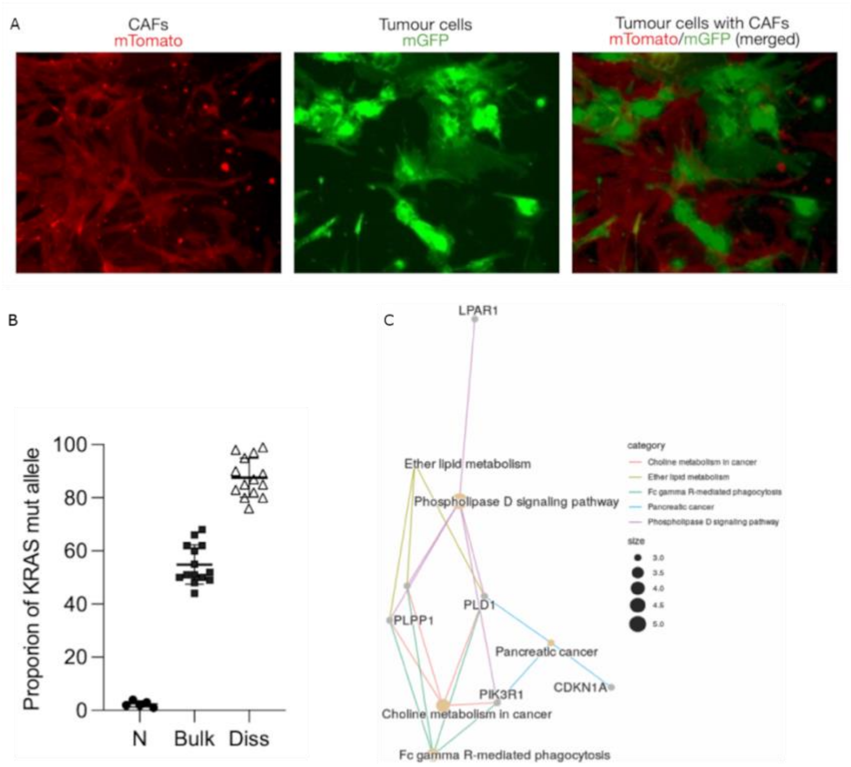


**Supplementary Figure S3 | Characterization of tumour cells, *KRAS* allelic imbalance, and pathway enrichment in porcine PDAC.**

**(A)** Fluorescence imaging of tumour-derived cells from KPC/R26$^{mT/mG}$ pigs. Cancer cells expressing mutant *KRAS*/*TP53* are labelled by mGFP (green), while cancer-associated fibroblasts (CAFs) are labelled by mTomato (red). The merged image demonstrates the coexistence of tumour cells and stromal components within the culture.

**(B)** Quantification of mutant *KRAS* allele frequency in normal pancreas (N), bulk tumour tissue (Bulk), and microdissected tumour epithelium (Diss). Microdissected samples show markedly increased mutant allele frequencies compared to bulk tissue, consistent with stromal dilution effects.

**(C)** Pathway enrichment analysis of differentially expressed genes in porcine PDAC, highlighting cancer-associated pathways including phospholipase D signalling, choline metabolism, ether lipid metabolism, and pancreatic cancer-related signalling networks. Node size reflects enrichment significance, and edges indicate gene–pathway associations.
