## Supplemental Data 1 for "A genetically engineered porcine model of pancreatic ductal adenocarcinoma captures the biological complexity of the human disease": Supplementary Tables 1 and 2.pdf

Supplementary Table S1. List of primers.

| Primer Name | Sequence | Application |
|---|---|---|
| EIF2B1 Forward | ATGTCCCCGTCACTGTGGTC | qPCR |
| EIF2B1 Reverse | CGCATACGGCCATCTGGTTG | qPCR |
| VIM Forward | GCTTCTTTGGCACGTCTTGA | qPCR |
| VIM Reverse | TGTTCCTGGATCTGAGCCTG | qPCR |
| ECDH Forward | TTATCCCTCCCATCAGCTGC | qPCR |
| ECDH Reverse | CTATCCAGAGGCTCCGTCAC | qPCR |
| αSMA Forward | AGCTTCGTGTTGCCCCAGAA | qPCR |
| αSMA Reverse | CAGAGGCGTAGAGGGACAGC | qPCR |
| CD44 Forward | CTGACCTCTGCAAAGCCTTC | qPCR |
| CD44 Reverse | CTGGGAGGTATTGGATGCGA | qPCR |
| S100A2 Forward | AGTACTCGGGCCAAGAAGGTG | qPCR |
| S100A2 Reverse | GAGGGCCAGGAAAACGGCAT | qPCR |
| KRAS-loxP Forward | AAAGCGGTACTTGCCTTTAAT | Cre recombination PCR |
| KRAS-loxP Reverse | TGAGGAAAAGAACAGTGCAAA | Cre recombination PCR |
| TP53-loxP Forward | TGAGGAATTTGTATGCCAAGG | Cre recombination PCR |
| TP53-loxP Reverse | TTCCACCAGTGAATCCACAA | Cre recombination PCR |
| KRAS AQ DNA For | GCCTGCTGAAAATGACTGA | Pyrosequencing |
| KRAS AQ DNA Rev | TGTACCCAAAGGATGGGCCT | Pyrosequencing |
| KRAS AQ DNA Seq | TGTGGTAGTTGGAGCTG | Pyrosequencing |
| KRAS AQ cDNA For | GCCTGCTGAAAATGACTGA | Pyrosequencing |
| KRAS AQ cDNA Rev | CATGTACTGGTCCCTCATT | Pyrosequencing |
| KRAS AQ cDNA Seq | TGTGGTAGTTGGAGCTG | Pyrosequencing |
| KRAS-cf Forward | ACACAGCAGGTCAAGAGGAG | EV qPCR |
| KRAS-cf Reverse | AACAAAGAAAGCCCTCCCCAG | EV qPCR |
| TP53-cf Forward | GGCGAGTATTTCACCCTCCA | EV qPCR |
| TP53-cf Reverse | AGCTCCAAGGCGTCATTCAG | EV qPCR |
| ACTB-cf Forward | CGGCATCCACGAAACTACCT | EV qPCR |
| ACTB-cf Reverse | CTCCTTCTGCATCCTGTCGG | EV qPCR |

Supplementary Table S2. List of primary and secondary antibodies. The application for which the antibody was used is indicated in brackets. In each line, the antibody is followed by the supplier and the corresponding catalogue number.

| | | |
|---|---|---|
| Cytokeratin 19 (IHC) | myBioSource | #MBS822263 |
| α-SMA (IHC) | Abcam | #ab150301 |
| Vimentin (IHC) | Cell Signaling | #5741 |
| E-cadherin (IHC) | Cell Signaling | #3195 |
| GATA6 (IHC) | Cell Signaling | #5851 |
| RAS(G12D) (FACS) | NewEast Biosciences | #26036 |
| Goat Anti-Rabbit IgG-HRP (IHC) | Santa Cruz | #sc-2780 |
| Giat Anti-Mouse IgG AF488 | Invitrogen | #A-11001 |
| CD3ε-PeCy7 (clone PPT3) (FACS) | BD Biosciences | #561477 |
| CD3ε-PE (clone BB23-8E6-8C8) (FACS) | BD Biosciences | #561485 |
| CD4-PerCPCy5.5 (clone 74-12-4) (FACS) | BD Biosciences | #561474 |
| CD8α-FITC (clone 76-2-11) (FACS) | BD Biosciences | #551303 |
| CD8β-Biotin (clone PG164A) (FACS) | Washington State University | #PG2020 |
| CD2-Biotin (clone MSA4) (FACS) | Kingfisher Biotech | #WS0590S-100 |
| CD25-AF647 (clone K231.3B2) (FACS) | BioRad | #MCA1736A647 |
| TCR1δ-Biotin (clone PGBL22A) (FACS) | Kingfisher Biotech | #WS061S-100 |
| IFN-γ-Biotin (clone CC302) (FACS) | BioRad | #MCA1783A647 |
| TNF-α-PECy7 (clone Mab11) (FACS) | Biolegend | #502926 |
| Perforin-BV421 (clone δG9) (FACS) | Biolegend | #308106 |
| FoxP3-PeCy7 (clone FJK-16s) | eBioscience | #E07638-1632 |
| Streptavidin-BV421 | BD Biosciences | #563259 |
| Streptavidin-BV711 | BD Biosciences | #563262 |
| Streptavidin-PE | BD Biosciences | #554061 |
